# Agricultural Pollution Risks Influence Microbial Ecology in Honghu Lake

**DOI:** 10.1101/244657

**Authors:** Maozhen Han, Melissa Dsouza, Chunyu Zhou, Hongjun Li, Junqian Zhang, Chaoyun Chen, Qi Yao, Chaofang Zhong, Hao Zhou, Jack A Gilbert, Zhi Wang, Kang Ning

**Author notes:** These authors contributed equally to this work (Maozhen Han, Melissa Dsouza, Chunyu Zhou).

## Abstract

**Background:** Agricultural activities, such as stock-farming, planting industry, and fish aquaculture, can influence the physicochemistry and biology of freshwater lakes. However, the extent to which these agricultural activities, especially those that result in eutrophication and antibiotic pollution, effect water and sediment-associated microbial ecology, remains unclear.

**Methods:** We performed a geospatial analysis of water and sediment associated microbial community structure, as well as physicochemical parameters and antibiotic pollution, across 18 sites in Honghu lake, which range from impacted to less-impacted by agricultural pollution. Furthermore, the co-occurrence network of water and sediment were built and compared accorded to the agricultural activities.

**Results:** Physicochemical properties including TN, TP, NO_3_^-^-N, and NO_2_^-^-N were correlated with microbial compositional differences in water samples. Likewise, in sediment samples, Sed-OM and Sed-TN correlated with microbial diversity. Oxytetracycline and tetracycline concentration described the majority of the variance in taxonomic and predicted functional diversity between impacted and less-impacted sites in water and sediment samples, respectively. Finally, the structure of microbial co-associations was influenced by the eutrophication and antibiotic pollution.

**Conclusion:** These analyses of the composition and structure of water and sediment microbial communities in anthropologically-impacted lakes are imperative for effective environmental pollution monitoring. Likewise, the exploration of the associations between environmental variables (e.g. physicochemical properties, and antibiotics) and community structure is important in the assessment of lake water quality and its ability to sustain agriculture. These results show agricultural practices can negatively influence not only the physicochemical properties, but also the biodiversity of microbial communities associated with the Honghu lake ecosystem. And these results provide compelling evidence that the microbial community can be used as a sentinel of eutrophication and antibiotics pollution risk associated with agricultural activity; and that proper monitoring of this environment is vital to maintain a sustainable environment in Honghu lake.

## Background

Water ecosystems, especially inland lakes, have suffered from eutrophication associated with increased agricultural activity comprising fish aquaculture as well as crop and livestock farming on surrounding lands(Brooks et al. 2016, Geist and Hawkins 2016, Williams et al. 2016). Improperly managed agricultural activities, such as excessive and/or improper use of fertilizers and/or pesticides, can cause eutrophication, which can negatively impact biodiversity(Williams et al. 2004). Previous studies have focused on the impact of this pollution on macro-organismal communities(Verdonschot et al. 2011, Williams et al. 2004) and in comparison, microbial ecology remains relatively understudied.

Agricultural pollution alters the physicochemical properties of water ecosystems(Baquero et al. 2008), which changes the associated microbial community composition and structure. In particular, nitrogen and phosphorus content, water temperature, and pH can fundamentally influence the microbiome(Bowles et al. 2014, Lindström et al. 2005, Xu et al. 2010). However, few studies have quantified the impact of organic pollutants such as herbicides and antibiotics. Determining the ecosystems resilience to such disturbance can aid conservation and help in the development of remediation strategies. There is an urgent need to develop sustainable approaches that establish a balanced relationship between the environment and agricultural production.

Antibiotics are widely utilized in livestock and fish aquaculture to promote animal growth and for the prophylactic or curative treatment of infectious disease(Lee et al. 2007), yet surface runoff of the introduction of treated sewage can introduce antibiotic pollution into local water bodies. Antibiotics inhibit microbial activity and can therefore influence biogeochemical processes in these ecosystems(Sengupta et al. 2013) and potentially select for antibiotic resistance mechanisms in environmental bacteria(Cherkasov et al. 2008). In addition, animal sewage can introduce animal-associated antibiotic resistant bacteria into these environments (Wang et al. 2017a), and as such it is necessary to have better quantification of the fitness and recovery rates of these resistant microbes upon release into the environment(Pei et al. 2006).

Honghu lake is a large, shallow eutrophic lake with an area of ~350 km^2^ and an average depth of ~1.5 m (**Figure 1**). It is located between the irrigation channel of the Four-lake main canal and the Yangzi River. Over the last five decades, Honghu lake has been extensively altered by flood regulation, irrigation, fish aquaculture, shipping, and water supply demands(Ban et al. 2014, Zhang 1998). Today, more than 40% of the lake area is used for large-scale aquaculture(Zhang et al. 2017). The intensive use of Honghu lake resources and the emission of sewage and other pollutants including fertilizers, pesticides, and antibiotics into the lake have led to a severe degradation of its water quality and an increase in the frequency of eutrophication events. In 2004, the Honghu Lake Wetland Protection and Restoration Demonstration Project(Zhang et al. 2017) was implemented to ameliorate the negative effects of severe water pollution, and one third of the lake area has been gradually protected under this provision. Consequently, Honghu lake represents a valuable, natural field site for investigating both the efficacy of the restoration program and the long-term effects of agricultural activities, such as the excessive application of antibiotics on water microbial communities.

**Figure 1.**
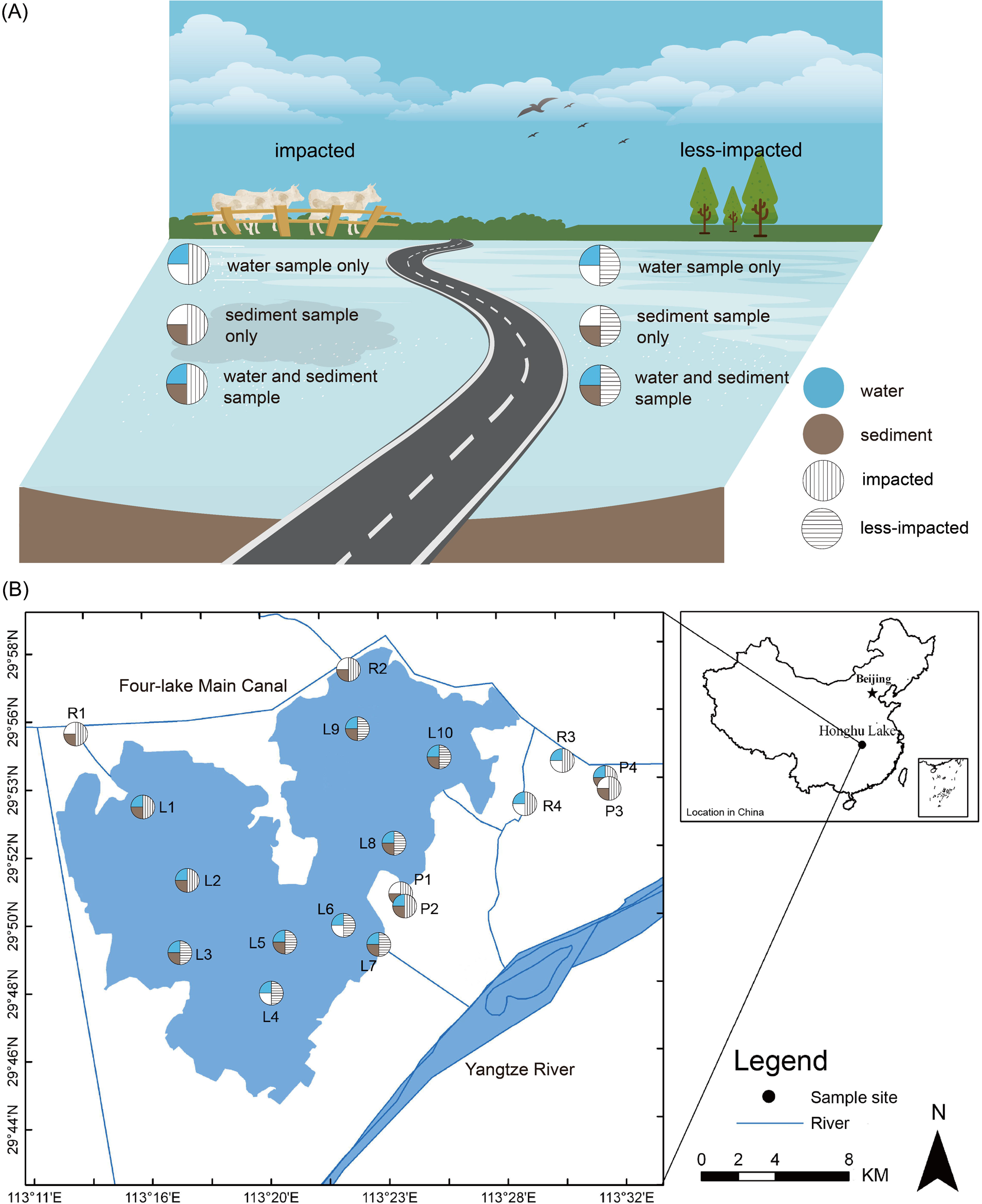
Geographic distribution of all sampling sites in Honghu lake. L: Lake; P: Pond; R: River. **(A)** Definitions of the various sampling strategies. **(B)** Locations of sampling sites and the distribution of the sampling medium collected at each site.

This study aimed to understand the geospatial influence of pollution on the water and sediment-associated microbial communities in Honghu lake. We performed 16S rRNA amplicon sequencing to characterize the microbial ecology, which was correlated against physicochemistry and antibiotic concentrations in these environments. This research was guided by the following scientific questions: (i) How does microbial diversity differ between water and sediment, and how is this influenced by the intensity of agricultural pollution? (ii) Which physicochemical properties and antibiotics are correlated with changes in microbial community structure? (iii) How are the co-occurrence relationships between microbiota influenced by the intensity of agricultural pollution (impacted and less-impacted)? Importantly, this baseline study aims to generate microbioal biomarkers of pollution, and to identify the ecological trends that could be used to provide a sentinel of pollution events in this lake environment.

## Results

### Physicochemical and antibiotic characterization

Physicochemical characteristics and antibiotic concentrations were determined for all water and sediment samples (**Figure 1, Supplementary Tables S1, S2, and S4**). Significant differences in pH (*t*-test, *P*=0.0037), ORP (*t*-test, *P*=0.00068), TN (*t*-test, *P*=0.035), and NH_4_^+^-N (*t*-test, *P*=0.045) were observed between water samples from impacted and less-impacted (control) sites (**Supplementary Table S1**). Samples from impacted sites were significantly more acidic and had greater concentrations of ORP, TN, and NH_4_^+^-N when compared to less-impacted sites (**Supplementary Table S1**). Similarly, sediment samples maintained significantly different Sed-OM (*t*-test, *P*=0.002), Sed-LP (*t*-test, *P*=0.0335), Sed-TN (*t*-test, *P*=0.0013), and Sed-TP (*t*-test, *P*=0.021) levels between impacted and less-impacted sites (**Supplementary Table S2**). Less-impacted sediment had greater concentrations of Sed-OM, Sed-LP, and Sed-TN when compared to the impacted sites (**Supplementary Table S2**), which may largely be due to the decomposition of plant material over the preceding winter months. Between impacted and less-impacted sites, the antibiotics ofloxacin (OFL, *t*-test, *P*=0.0079) and sulfamethoxazole (SMZ, *t*-test, *P*=0.043) had significantly different concentrations in water samples, while sulfamerazine (SMR, t-test, *P*=0.021) was significantly different in sediment samples (**Supplementary Table S4**); in both cases concentrations were greater in impacted sites.

### Microbial diversity and community structure

A total of 28 water and sediment samples generated 4,441,405 paired-end 16S rRNA reads, which clustered into 7,785 OTUs (**Supplementary Information**). Microbial alpha diversity was significantly greater in sediment samples (Chao1 (*t*-test, *P*=0.0045, **Supplementary Table S5**) and PD whole tree (*t*-test, *P*=0.003, **Supplementary Table S5))**. The microbial alpha diversity in sediment samples was significantly different between impacted and less-impacted sites (*t*-test, *P*=0.0445, **Supplementary Table S5**). However, no significant difference in alpha diversity was observed in water samples.

A total of 53 microbial phyla were identified across all samples (**Figure 2A**), and were differentiated between water and sediment samples (**Figure 2B**), and between impacted and less-impacted sites (PERMANOVA, Bray-Curtis distance, *P* <0.01). In water samples, Proteobacteria (*t*-test, *P* <0.05), Cyanobacteria (*t*-test, *P* <0.05), and Gemmatimonadetes (*t*-test, *P* <0.05) were significantly different between impacted and less-impacted sites (**Figure 2C**). While in sediment samples, Actinobacteria (*t*-test, *P* <0.01), Firmicutes (*t*-test, *P* <0.05), Bacteroidetes (*t*-test, *P* <0.05), Nitrospirae (*t*-test, *P* <0.05), and OP8 (*t*-test, *P* <0.05) were significantly different between impacted and less-impacted sites (**Figure 2D**).

**Figure 2.**
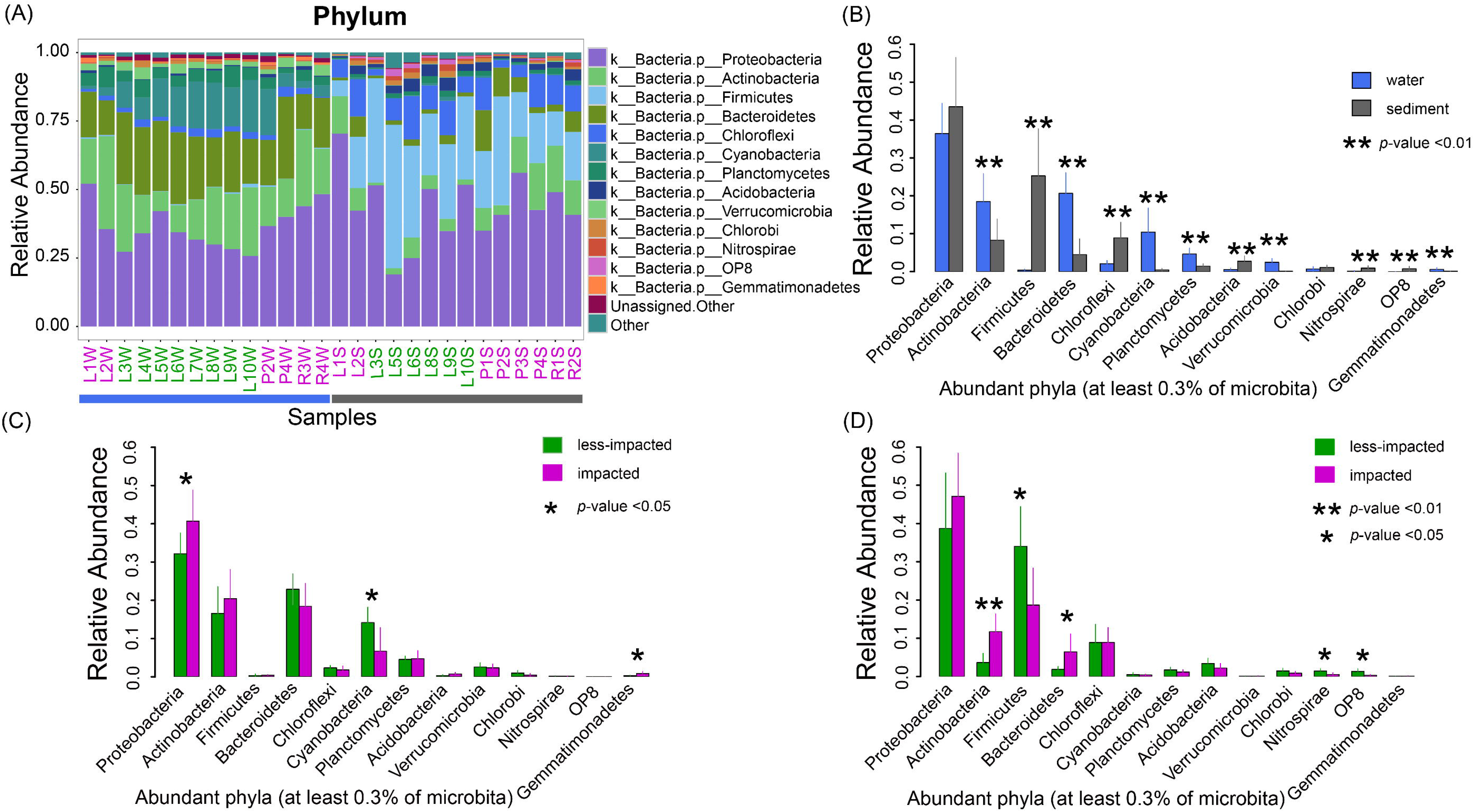
Taxonomic composition and relative abundances of microbial taxa in water and sediment samples. **(A)** Taxonomic composition of each sample at the phylum level. ‘Other’ represents all phyla not in the top 13 phyla. **(B)** Bar plot highlighting differences between water and sediment samples at phylum level. ** represents *p*-values <0.01 and determined by a student *t*-test. **(C)** Bar plot highlighting differences in water samples at the phylum level between impacted and less-impacted groups. * represents *p*-values <0.05 and determined by a student *t*-test. **(D)** Bar plot highlighting differences in sediment samples at the phylum level between impacted and less-impacted groups. * represents *p*-values <0.05 and determined by a student *t*-test. ** represents *p*-values <0.01 and determined by a student *t*-test.

Core-OTUs were defined as a set of OTUs that were identified in all samples analyzed, and Pan-OTUs were defined as a set of OTUs that were identified in at least one sample. Core- and Pan-OTUs were determined for all water and sediment samples (**Figure 3**). A total of 132 Core-OTUs and 7,418 Pan-OTUs were identified in less-impacted sites, while impacted sites maintained 201 Core-OTUs and 7,706 Pan-OTUs (**Supplementary Figure S3 and Supplementary Figure S4**). The Core-OTUs from both the impacted and less-impacted sites were dominated by Proteobacteria, specifically *Janthinobacterium* (**Supplementary dataset 4 and 5**), while Acidobacteria were enriched at the impacted sites (2.79%±1.30%, **Supplementary dataset 4**).

**Figure 3.**
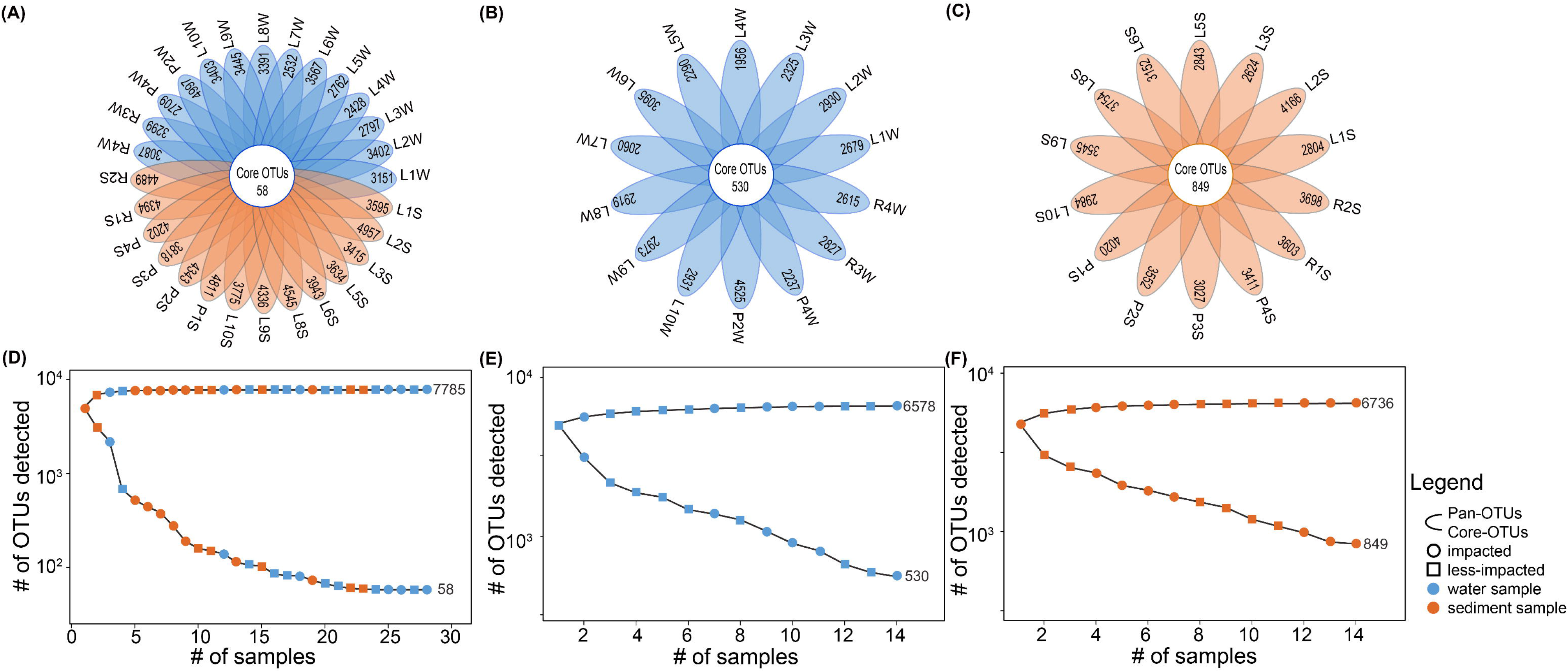
Core- and Pan-OTUs of water and sediment samples from Honghu lake. Flower plots showing the number of sample-specific OTUs (in the petals) and Core-OTUs (in the center) for **(A)** all samples, **(B)** all water samples, and **(C)** all sediment samples. OTU accumulation curves for Pan-OTUs (upper) and Core-OTUs (lower) for **(D)** all samples, **(E)** all water samples, and **(F)** all sediment samples from Honghu lake.

Microbial beta diversity was further assessed by UPGMA clustering using the unweighted UNiFrac distance matrix. We observed clustering by sampling medium (**Figure 4A** and **Supplementary Figure S5**) and by level of agricultural activity within water and sediment samples (**Figure 4B**). Importantly, greater differences in beta diversity were observed between impacted and less-impacted sites in sediment samples as compared to water samples (**Figure 4B and 4C)**.

**Figure 4.**
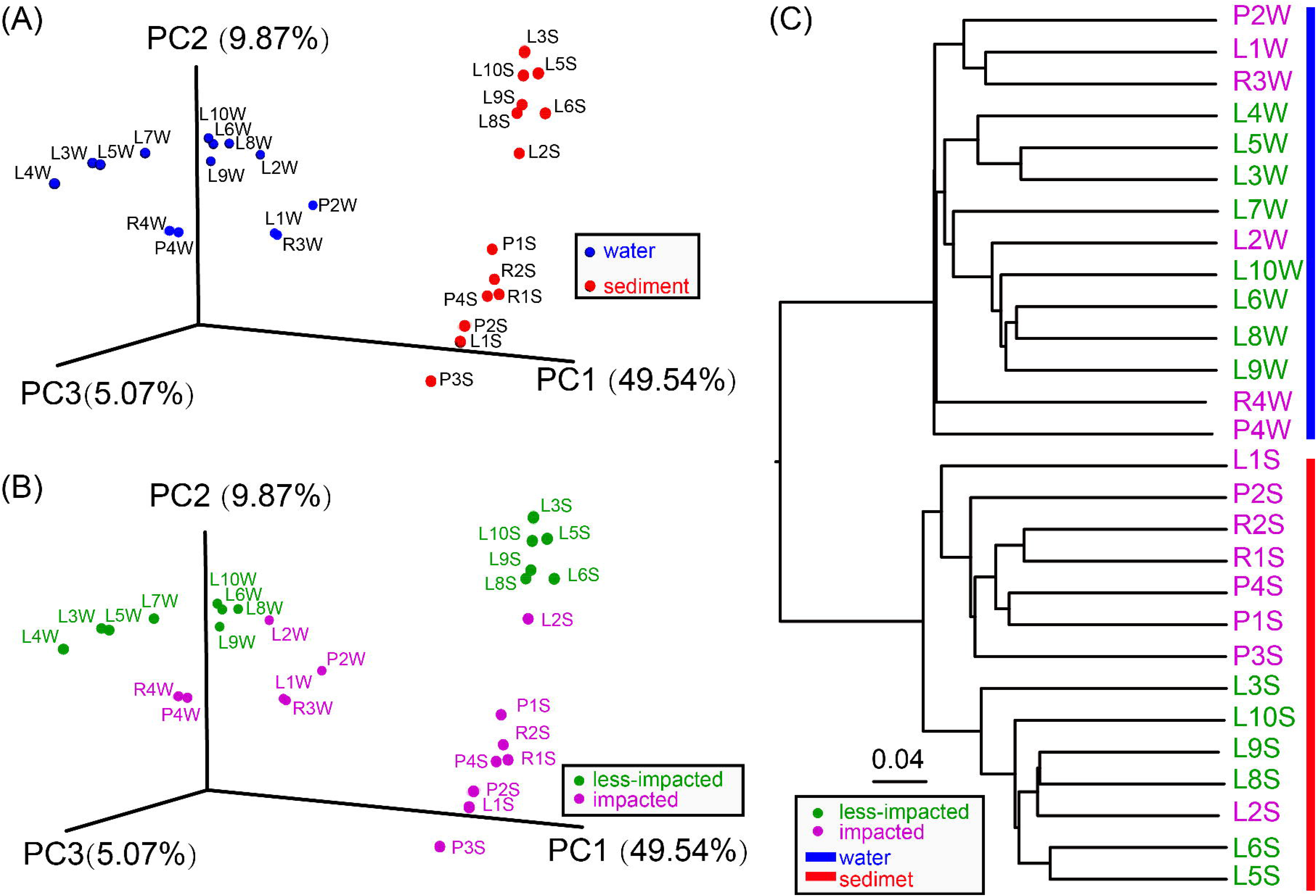
PCoA plots and UPGMA-based clustering of water and sediment microbial communities. Unweighted UNitrac dissimilarity matrix scores for all samples visualized in a PCoA plot to demonstrate the dissimilarity of the microbial community structure between samples by **(A)** sampling medium (water vs. sediment) and by **(B)** impacted and less-impacted groups. **(C)** UPGMA-based clustering tree of microbial communities using an unweighted UNitrac distance matrix. The green and pink font represent less-impacted and impacted groups, respectively. The blue and red bars mark water and sediment samples, respectively.

### Functional properties predicted by PICRUSt

We observed clustering of water and sediment microbial communities based on the relative abundance of their predicted functional profiles (**Supplementary Figure S6**) (PERMANOVA, Bray-Curtis distance, *P* <0.0001). In water samples, functional groups including amino acid related enzymes, peptidases, oxidative phosphorylation, purine metabolism, pyrimidine metabolism, DNA repair and recombination proteins, and arginine and proline metabolism were enriched (**Supplementary Figure S7**). Likwise, in sediment samples we observed an enrichment of functional groups including ribosome biogenesis, secretion system, two-component system, ABC transporters, and pyruvate metaolism (**Supplementary Figure S7**). When investigating agricultural pollution risks, we observed significant differences in the relative abundances of the predicted functional profiles between impacted and less-impacted groups of water samples (PERMANOVA, Bray-Curtis distance, *P* <0.05). For these samples, the relative abundances of DNA repair and recombination proteins (*t*-test, *P* <0.05), purine metabolism (*t*-test, *P* <0.05), secretion systems (*t*-test, *P* <0.05), oxidative phosphorylation (*t*-test, *P* <0.05), pyrimidine metabolism (*t*-test, *P* <0.05), amino acid related enzymes (*t*-test, *P* <0.05), and arginine and proline metabolism (*t*-test, *P* <0.05) were significantly different between impacted and less-impacted sites (**Supplementary Figure S7**). In contrast, we observed no significant differences in sediment functional profiles between impacted and less-impacted sites.

### Correlating physicochemical properties with Microbial diversity

Physicochemical properties including NH_4_^+^-N, TN, ORP, TP, Tur, COD_Mn_, and Chl (**Supplementary dataset 6**) were significant explanatory factors that determined the observed clustering pattern of the water microbial communities at impacted sites (**Figure 5A, Supplementary Figure S8A**), while pH and DO determined the water microbial community structure at less-impacted sites (**Figure 5A, Supplementary Figure S8A**). For sediment samples, Sed-LP, Sed-TN, and Sed-OM (**Supplementary dataset 7**) were identified as significant explanatory factors shaping the observed clustering pattern at less-impacted sites and Sed-TP for impacted sites (**Figure 5B, Supplementary Figure S8B**). Based on distance correlations and the statistical significance of Mantel’s r-statistic, water physciochemical properties including TN, ORP, NO_3_^-^-N, and NO_2_^-^-N, were strongly correlated with taxonomic and functional composition (**Figure 6A**). For sediment samples, Sed-OM and Sed-TN were strongly correlated with taxonomic data (**Figure 6B**).

**Figure 5.**
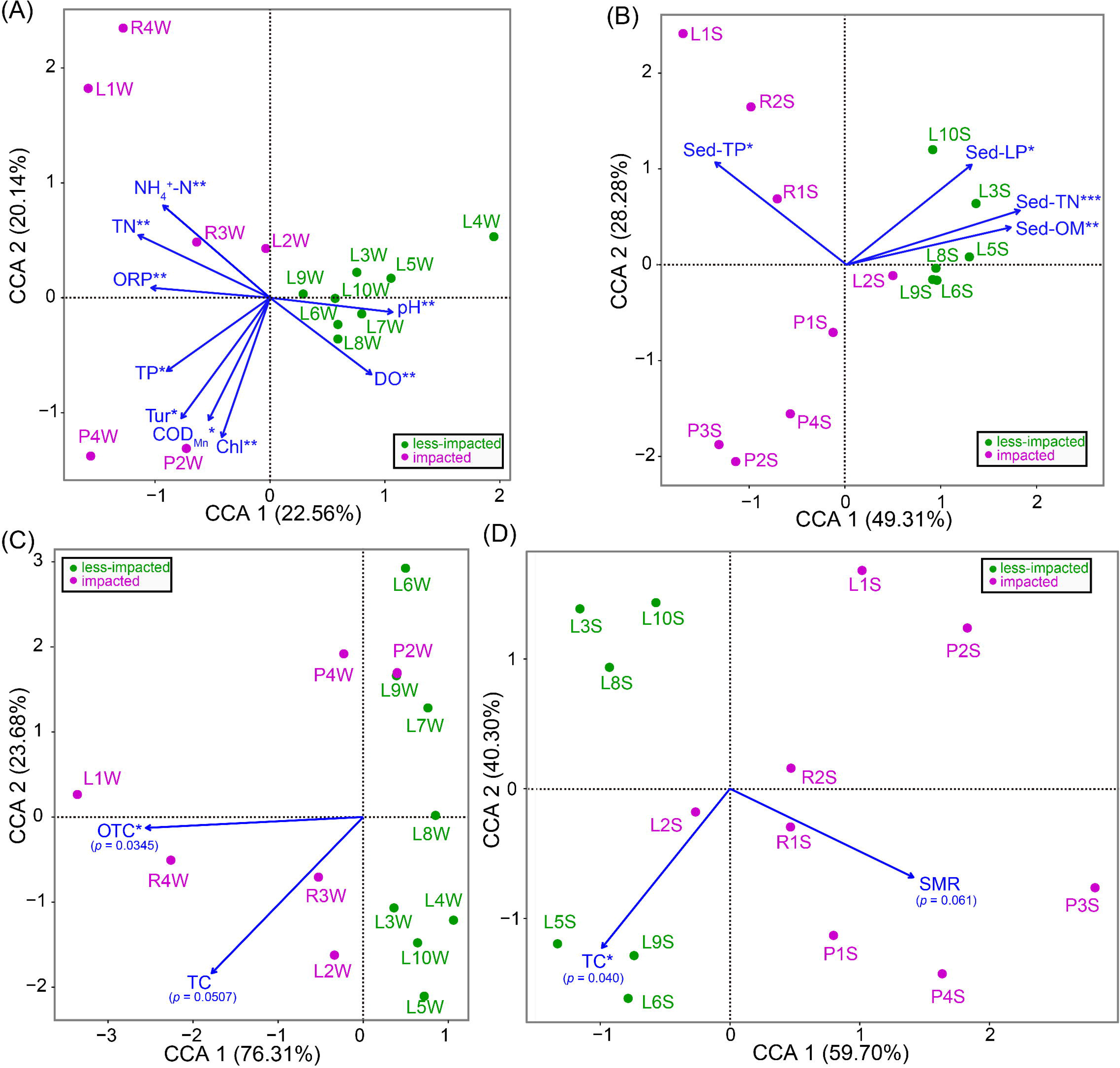
Canonical correspondence analysis plots of physicochemical properties and antibiotic data driving water and sediment microbial community structure. Physicochemical properties of **(A)** water samples **(B)** and sediment samples from Honghu lake, and antibiotic data for **(C)** water samples and **(D)** sediment samples from Honghu lake. *** represents *p*-values <0.001, ** represents *p*-values <0.01 and * represents *p*-values <0.05.

The antibiotic oxytetracycline (OTC) was the primarily explanatory factor for water microbial diversity variance at impacted sites (**Figure 5C, Supplementary Figure S8A**). While in sediment samples, sulfamerazine (SMR) was the primary factor responsible for the observed clustering of samples, including R1S, R2S, P1S, P3S, and P4S, from impacted sites (**Figure 5D, Supplementary Figure S8B**). Mantel correlation assessments were also performed between antibiotic data and compositional data for water and sediment samples (**Figure 6C, Figure 6D**, and **Supplementary Figure S9**). The oxytetracycline (OTC) antibiotic class was strongly correlated with water taxonomic and functional composition data (**Figure 6C**), while tetracycline (TC) was strongly correlated with sediment taxonomic and functional composition data (**Figure 6D**). The ofloxacin (OFL) antibiotic class was strongly correlated with taxonomic and functional composition data in sediment samples collected from less-impacted (control) sites (**Supplementary Figure S10B**). Additionally, COD_Mn_ and ciprofloxacin (CIP) were strongly correlated with taxonomic and functional composition in water samples collected from impacted sites (**Supplementary Figure S10C**).

**Figure 6.**
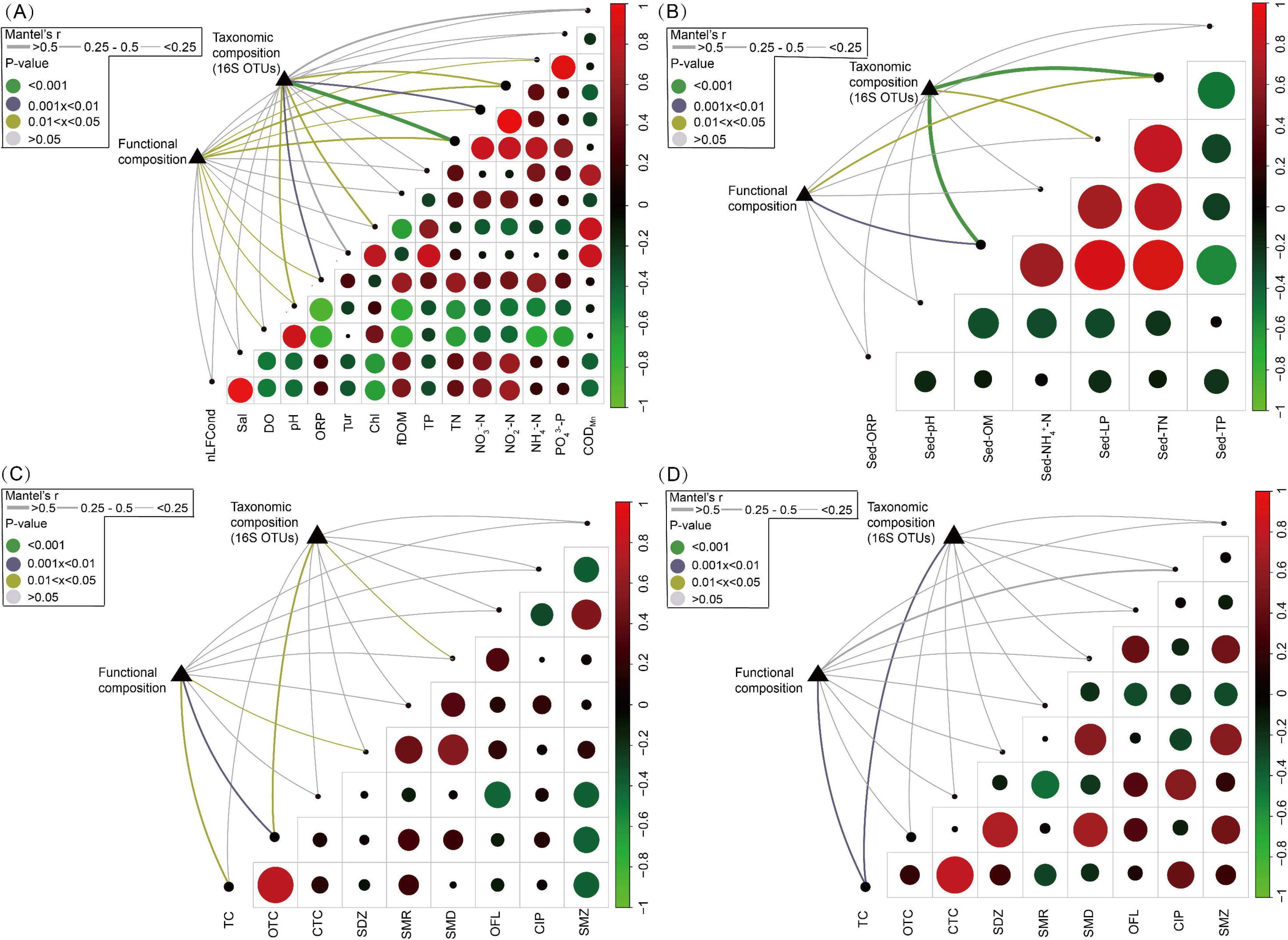
Environmental drivers of microbial community composition in water and sediment samples. Pairwise comparisons of **(A)** water and **(B)** sediment physicochemical properties with taxonomic and functional composition data. Color gradient represents Pearson’s Correlations Coefficients. Pairwise comparisons of **(C)** water and **(D)** sediment antibiotic concentration data with taxonomic and functional composition data. In all figures, the varying circle size represents the absolute value of the Pearson Correlation Coefficient between the two factors, the bar along the y-axis represents the value of the Pearson Correlation Coefficients, and the edge width represents the Mantel’s *r* statistic value for distance correlations and the edge color denotes the statistical significance based on 9,999 permutations. Abbreviations of physicochemical properties and antibiotics are listed in sub-section ***‘Physicochemical characterization and antibiotic analysis’***.

Moreover, we observed strong correlations between several OTUs, physicochemical properties, and antibiotic concentrations (**Supplementary Information**). In water samples, *Bacillus flexus* (denovo 71031, **Supplementary dataset 8**) was strongly correlated with TN (*r*=0.8675, fdr-*P*=7.89E-5, **Supplementary Figure S11C**), NH_4_^+^-N (*r*=0.8958, fdr-*P*=7.89E-5, **Supplementary Figure S11D)**, PO_4_^3-^-P (*r*=0.832, fdr-*P*=2.58E-4, **Supplementary Figure S11E**), and oxytetracycline (OTC, *r*=0.8381, fdr-*P*=3.62E-4, **Supplementary Figure S12B**).

### Biomarker discovery

In water samples, the LEfSe analysis identified 13 biomarkers for impacted sites and 12 for less-impacted sites. The most differentially abundant bacteria from impacted sites belonged to the phylum Proteobacteria, class Betaproteobacteria and class Gammaproteobacteria (**Figure 7A and 7B)**. These included members of the orders *Methylophilales, Nitrosomonadales*, and *Rhodocyclales* (**Figure 7A and 7)**. *Methylophilales* are known for their ability to metabolize methane under aerobic and microaerobic conditions(Beck et al. 2013) and *Nitrosomonadales* are significantly enriched in soils containing high concentrations of N fertilizer(Chávez-Romero et al. 2016). Water samples from less-impacted sites were overrepresented by *Oscillatoriophycideae* and *Synechococcophycideae* in *Cyanobacteria*; and *Saprospiraceae* in *Bacteroidetes* (**Figure 7A and 7B)**.

**Figure 7.**
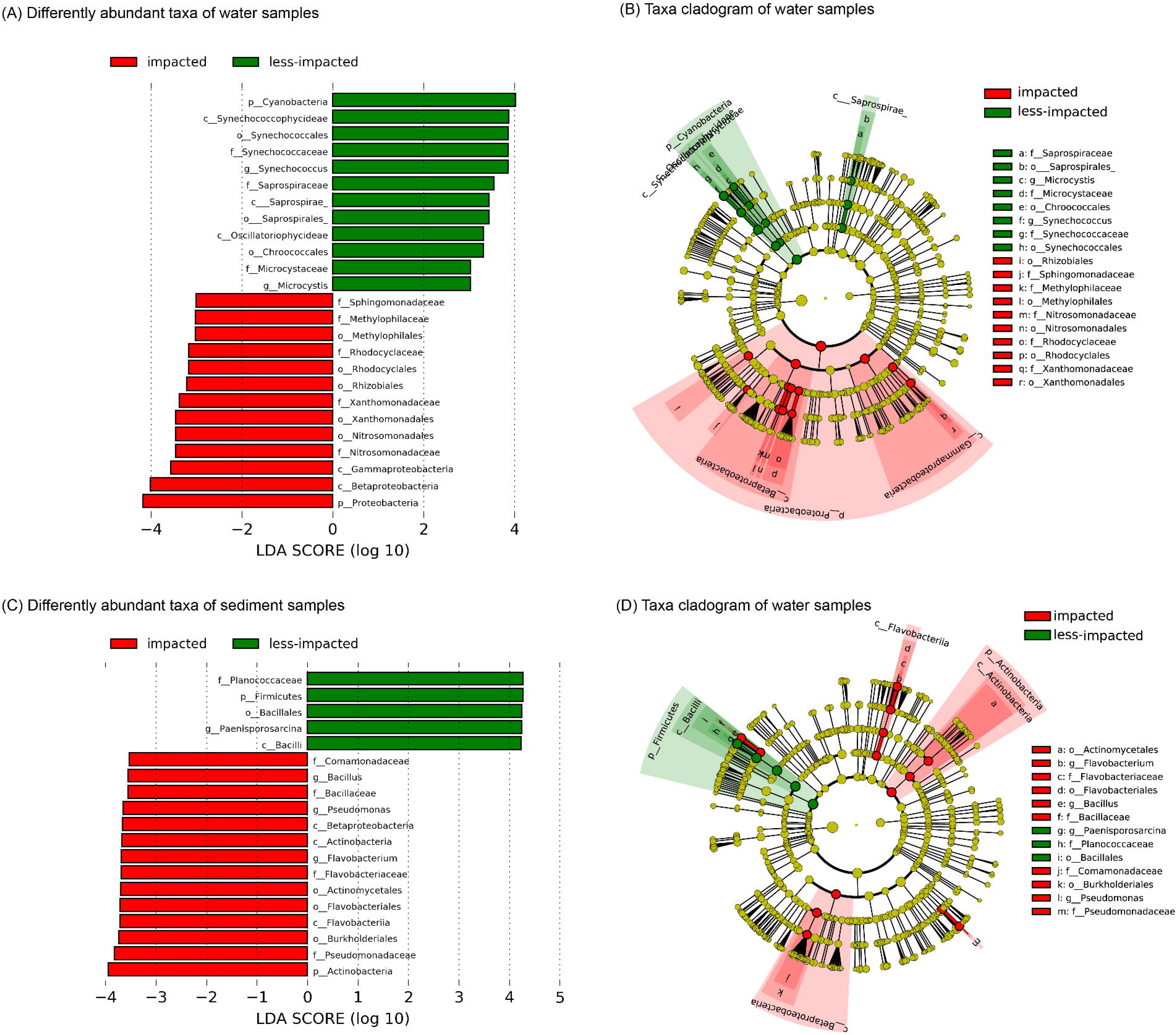
Biomarkers analysis of water and sediment microbial communities from impacted and less-impacted (control) sites. **(A)** Differentially abundant taxa of water samples; **(B)** Cladogram showing the phylogenetic structure of the microbiota from water samples; **(C)** Differentially abundant taxa of sediment samples; **(D)** Cladogram showing the phylogenetic structure of the microbiota from sediment samples.

In sediment samples, the LEfSe analysis reported 14 biomarkers enriched in impacted sites and 5 enriched in less-impacted sites (**Figure 7C and 7D)**. Biomarkers in samples from impacted sites mainly comprised members of the phylum Actinobacteria, family *Pseudomonadaceae*, order *Burkholderiales*, and *class Flavobacteriia*. For sediment samples from less-impacted sites, bacteria that were differentially abundant include members of *Paenisporosarcina* genus and candidate family planococcaceae, phylum Firmicutes, order Bacillales, and class Bacilli (**Figure 7C and 7D)**.

### Co-occurrence Network Analysis

Co-occurrence network analysis was performed to visualize and characterize co-occurrence patterns among members of water and sediment microbial communities. The water and sediment network comprised 427 nodes and 189 edges (**Figure 8A**) and 443 nodes and 2,877 edges (**Figure 8B**), respectively. The density of the water and sediment network was 0.002 and 0.023, respectively. These results suggest that the sediment microbial network was more connected than the water network. Both networks exhibited a scale-free degree distribution pattern, whereby most OTUs had low degree values and fewer hub nodes had high degree values (**Supplementary Figure S15**).

**Figure 8.**
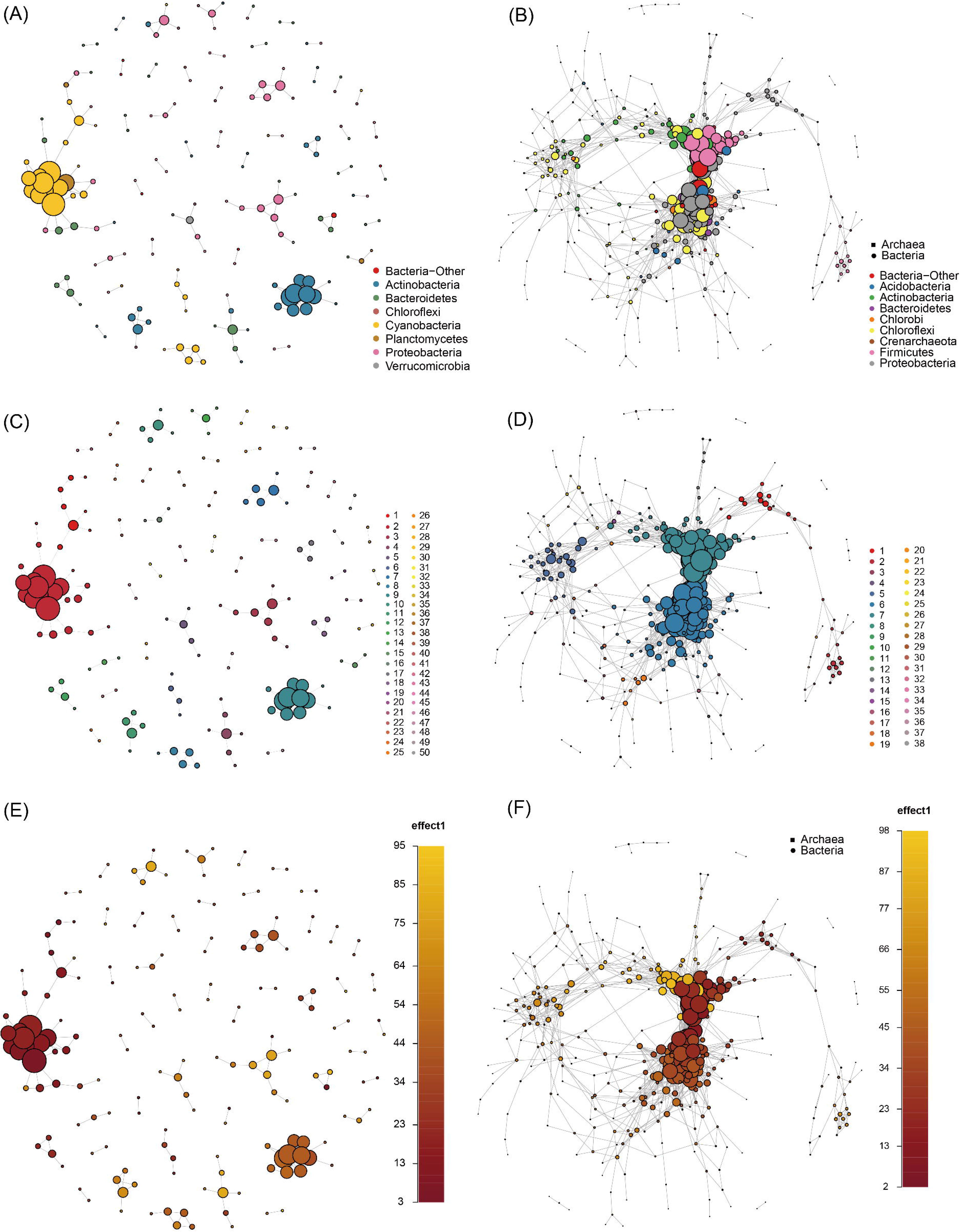
Co-occurrence network interactions of Honghu lake microbes in water and sediment samples. Network nodes represent OTUs and edges are represented as interactions between OTUs. All edges represent positive, strong (Spearman’s ρ>0.8), and significant (*p*-value<0.001) correlations. The size of each node is proportional to the node degree. Networks of **(A)** water and **(B)** sediment samples displaying co-occurrence patterns of OTUs grouped at the phylum level. Modules were identified using the WalkTrap community detection algorithm in **(C)** water and **(D)** sediment samples. Water **(E)** and **(F)** sediment networks investigating the effect of long-term agricultural activities wherein each node was colored as function of its relative abundance in impacted and less-impacted sites.

We detected modules in water and sediment networks using the WalkTrap community detection algorithm. The modularity of the water and sediment network was 0.878 and 0.559, respectively. A total of 50 clusters with the largest membership of 22 was observed for the water network (**Figure 8C**). Likewise, for the sediment network we observed a total of 38 clusters with the largest membership of 111 (**Figure 8D**). In the sediment network, most OTUs in module 6 (111 nodes) were members of Anaerolineae of the phylum Chloroflexi and Beta-, Delta-, and Gammaproteobacteria. Additionally, most OTUs in module 7 (81 nodes) of the sediment network were members of the Planococcaceae, a family within the order Bacillales. When compared to the sediment network, we observed fewer and smaller hubs in the water network. In this network, most OTUs in module 2 (22 nodes) and module 9 (12 nodes) were members of the genus *Synechococcus* within the order Synechococcales and ACK-M1 within the order Actinomycetales, respectively.

We also examined the effect of prolonged agricultural activities on the co-occurrence patterns of water and sediment microbial communities. For this, each node in the water and sediment network was colored as a function of its relative abundance across samples from impacted and less-impacted (control) sites (**Figure 8E and 8F**). In both networks, we observed higher connectedness among OTUs associated with less-impacted samples as compared to those associated with samples from impacted sites. This observation was confirmed when we generated sub-networks for impacted and less-impacted sediment samples by selecting OTUs associated with these samples and all edges among them from the overall sediment co-occurrence network (**Supplementary Figure S16**). We observed higher connectedness in microbes associated with less-impacted samples (measured as node degree, 3.746) as compared to those associated with samples from impacted sites (1.397).

## Discussions

The extensive application of chemical compounds such as fertilizers, herbicides, and antibiotics, can profoundly influence the cycling and accumulation of nutrients in the sediment and water column of Honghu lake(Chen et al. 2008). These agricultural practices can negatively impact not only the physicochemical properties, but also the biodiversity of microbial communities associated with the lake ecosystem(Baquero et al. 2008). These changes in microbial community composition can in turn affect nutrient cycling and organic matter decomposition, thus impacting overall agricultural productivity.

In our study, we analyzed water and sediment samples from Honghu lake, assessing its microbiome, physicochemical properties, and antibiotic concentrations. We found that despite low human activity, high concentrations of Sed-LP, Sed-TN, and Sed-OM were observed at less-impacted (control) sites, probably due to the abundance of submerged plants. We speculate that the decay of these plants during winter subtantially increases organic matter, total nitrogen (Sand-jensen 1998), and total phosphorus(Horppila and Nurminen 2003) in sediment samples. Hence, as expected from previous research, we found that both water and sediment microbial community structure was correlated with TP and TN concentration(Han et al. 2016, Takamura N 2003). Moreover, in water samples, we observed that *Bacillus flexus* was strongly correlated with TN, NH_4_^+^-N, PO_4_^3-^-P, and oxytetracycline. More important, previous work on *Bacillus flexus* have shown that members of this species can degrade organic(Guo et al. 2013) and inorganic(Divyashree et al. 2009) nitrogen, thus making it a possible candidate for bioremediation in alkaline wastewater(Wang and Zhao 2013). Some strains of *B. flexus* also demonstrate strong phosphorus solubilization activity(Gechemba et al. 2015), and others demonstrated resistance to OTC(Sundaramanickam et al. 2015).

As to biomarkers in sediment samples from impacted sites, these included members of the *Hydrogenophaga* genus, belonging to *Burkholderiales* (Class *Betaproteobacteria)*, which have been previously associated with agricultural activities(Babujia et al. 2016). Moreover, members of the genus *Pseudomonas*, belonging to family *Pseudomonadaceae*, can play an important role in agricultural ecosystems, particularly those associated with plant growth-promotion and disease suppression were mentioned(Pesaro and Widmer 2006).

Co-occurrence network analysis showed that Anaerolineae forms a large component of microbial communities associated with sludge wastewater treatment plants wherein they may play important roles in organic degradation(Nielsen et al. 2009). The phylum Proteobacteria are known to easily metabolize soluble organic substrates(Shao et al. 2013). Among these classes, Deltaproteobacteria, a dominant group often observed in a variety of sediment samples, play an important role in degrading organic compounds to carbon dioxide(Ye et al. 2009). Members of *Synechococcus* are a cosmopolitan cyanobacterium often associated with toxic algal blooms and microcystin production(O’neil et al. 2012, Wood et al. 2017). Likewise, members of ACK-M1, in a recent study, exhibited chemotaxis towards ammonium in a water ecosystem, thus influencing nutrient cycling processes and microbial competitive interactions within this ecosystem(Dennis et al. 2013). The presence of these microbial taxa is indicative of the long-term effect of eutrophication in water environments.

## Conclusion

We analyzed the impacted sites and less-impacted sites of water and sediment samples from Honghu lake and surrounding river and pond sites. The microbiome was analyzed in the context of variable physicochemical properties and antibiotic concentrations. There were significant differences between impacted and less-impacted (control) groups in both water and sediment samples. These differences were observed in physicochemical properties, antibiotic concentration levels, and taxonomic structure. Physicochemical properties including TN, TP, NO3–N, and NO2–N were the main factors driving compositional differences in water samples. Likewise, in sediment samples, Sed-OM and Sed-TN were the main factors driving differences in taxonomic composition. The antibiotics, oxytetracycline and tetracycline were identified as the main drivers of taxonomic and functional structure in water and sediment samples, respectively. As for differences between impacted and less-impacted samples, we identified 25 biomarkers within water communities and 19 within sediment microbial communities. Finally, the co-occurrence network analysis revealed differences in co-occurrence patterns by sampling medium (water vs. sediment microbial communities) and by level of agricultural activity (impacted vs. less-impacted microbial communities). These results suggest that continued analyses of the composition and structure of water and sediment microbial communities in such anthropologically-impacted lake environments may provide valuable biomarker data to track pollution. The Honghu Lake Wetland Protection and Restoration Demonstration Project provided preliminary data that highlights the importance of monitoring biodiversity in water micro-ecosystems. Our present work allows further investigation into the impact of agricultural practices on water ecosystems and more importantly, into our ability to remediate these important ecosystems.

## Materials and Methods

### Sample sites and sampling processes

To investigate the differences in microbial community structure resulting from a wide range of anthropogenic activities, a total of 14 water samples and 14 sediment samples were collected from Honghu lake and surrounding rivers and ponds during 10-11 November 2015. Among these sites, L1 is the inlet of inflowing river, and sites L3, L8, L9, and L10 are relatively close to aquaculture areas. Meanwhile, to evaluate the primary source of the antibiotics of Honghu lake, the waters from four major connecting rivers (i.e., R1 to R4) of Honghu lake and four typical aquaculture ponds (i.e., P1 to P4), which can exchange water with Honghu lake, were collected(Wang et al. 2017b). In keeping with the Government Protection Zone definition(Zhang et al. 2017) and in taking into account the different sources of pollution at each site(Wang et al. 2017b) (treated sewage, crop, livestock, and fish aquaculture), all sampling sites were categorized into two groups—namely, the impacted and the less-impacted, control group(Zhang et al. 2017). Sampling sites labeled L1, L2, P1, P2, P3, P4, R1, R2, R3, and R4 were classified as impacted, while sites labeled L3, L4, L5, L6, L7, L8, L9, and L10 were classified as less-impacted (**Figure 1**).

For water sampling, 2 L of water at a depth of 0.3-0.5 m were collected at each site using a cylinder sampler. Approximately 1.5 L of sample was used for physicochemical characterization and antibiotic analysis. The remaining 500 mL of sample was size-fractionated using a 20 μm tulle and a 0.22 μm diameter pore size filter membrane (Tianjin Jinteng Experiment Equipment Co., Ltd). Microbial biomass was collected on 0.22 μm diameter pore size filter membranes. These membrane samples were stored onsite in a portable cooler with ice bags, then transported to the laboratory and stored at −80°C until DNA extraction. For sediment sampling, ~200 g of sediment (0-10 cm) was collected at each site and stored in a portable cooler with ice bags until its transportation to the laboratory for subsequent downstream analyses. Approximately 50 g of sediment was used for physicochemical characterization and antibiotic analysis, whilst the remainder was dried in an Ultra-low Freeze Dryer (Christ, German) until no further weight changes were observed. The dried sediment (0.5 g) was used for DNA extraction.

### Physicochemical characterization and antibiotic analysis

#### Physicochemical characterization

Physicochemical data were collected for all water and sediment samples (**Supplementary Table S1 and Supplementary Table S2**). Physicochemical properties including water temperature (T), pH, dissolved oxygen (DO), temperature compensated conductivity (nlF Cond), salinity (Sal), oxidation-reduction potential (ORP), turbidity (Tur), chlorophyll-a (Chl-a), and fluorescent dissolved organic matter (fDOM) were measured for all water samples *in situ* by EXO2 (YSI, USA). Additional physicochemical properties including total phosphorus (TP), orthophosphate (PO_4_^3-^-P), total nitrogen (TN), ammonium nitrogen (NH_4_^+^-N), nitrate nitrogen (NO_3_^-^-N), nitrite nitrogen (NO_2_^-^-N), and potassium permanganate index (oxygen consumption, COD_Mn_) were assayed as described in previous work (Federation and Association 2005). For sediment samples, ORP (Sed-ORP) and pH (Sed-pH) were determined using a pH/ORP portable meter (YSI, USA). Sediment organic matter (Sed-OM) was determined in a muffle furnace at 550°C (Federation and Association 2005). Labile phosphorus (LP), NH_4_^+^-N (Sed-NH_4_^+^-N), total phosphorus (Sed-TP), and total nitrogen (Sed-TN) were measured by the NH_4_Cl extraction method, the KCl extraction method, the perchloric acid and sulfuric acid digestion method, and the Kjeldahl method, respectively(Bao 2005).

#### Antibiotic analysis

Based on a report of antibiotic usage in China(Yang et al. 2016), a total of 13 antibiotics were selected for detection in water and sediment samples (**Supplementary Table S4**). These antibiotics can be classified into three groups namely: (i) sulfonamides (SAs), including sulfadiazine (SDZ), sulfamerazine (SMR), sulfamate (SFM), sulfadimidine (SMD), sulfamonomethoxine (SMM), and sulfamethoxazole (SMZ); (ii) fluoroquinolones (FQs), including fleroxacin (FLE), ofloxacin (OFL), ciprofloxacin (CIP), and difloxacin (DIF); and (iii) the tetracycline group (TCs), including tetracycline (TC), oxytetracycline (OTC), and chlortetracycline (CTC). We determined the concentration of these antibiotics in water and sediment samples using a 2695 Waters Alliance system (Milford, MA, USA) equipped with an auto sampler-controlled binary gradient system, a micro vacuum degasser, and a 2998 Photodiode Array (PDA) detector. A detailed protocol of the antibiotic extraction process is described in **Supplementary Information**. Of the 13 antibiotics that were quantified, nine antibiotics including TC, OTC, CTC, SDZ, SMR, SMD, OFL, CIP, and SMZ were selected for further analysis in this study.

### DNA extraction and 16S rRNA gene sequencing

DNA was extracted from water filter membranes and dried sediment using a modified hexadecyltrimethylammonium bromide (CTAB) method(Cheng et al. 2014a, Cheng et al. 2014b, Porebski et al. 1997) (**Supplementary Information)**. All extracted DNA was dissolved in TE buffer and stored at –20°C until further use.

DNA samples were quantified using a Qubit^®^ 2.0 Fluorometer (Invitrogen, Carlsbad, CA) and DNA quality was assessed on 0.8% agarose gels. Approximately 5-50 ng of DNA was used as template for amplifying the V4-V5 hypervariable region of the 16S rRNA gene of microbiota for each sample. Sequences for the forward and reverse primers are "GTGYCAGCMGCCGCGGTAA" and "CTTGTGCGGKCCCCCGYCAATTC", respectively(Han et al. 2016). The sequencing library was constructed using a MetaVx™ Library Preparation kit (GENEWIZ, Inc., South Plainfield, and NJ, USA). Indexed adapters were added to the ends of the 16S rDNA amplicons by limited cycle PCR. DNA libraries were verified using an Agilent 2100 Bioanalyzer (Agilent Technologies, Palo Alto, CA, USA) and quantified by Qubit^®^ 2.0 and quantitative PCR (Applied Biosystems, Carlsbad, CA, USA). All sequencing reactions were performed on the Illumina MiSeq platform using paired-end sequencing technology (2*300 bp).

### Quality control, OTU clustering, and taxonomy assignment

All 16S rRNA gene amplicons were processed according to the ensuing criteria and sequences below the set quality threshold were excluded from subsequent analyses. Firstly, paired-end reads were spliced using the ‘make.contigs’ command in mothur(Schloss et al. 2009) (version 1.25.0) with default settings. All reads containing ambiguous base calls (N), those longer than 500 bp, and those shorter than 300 bp were removed. Putative chimeras were identified using the SILVA database(Quast et al. 2013) (Release 123) and removed using the ‘chimera.uchime’ and ‘remove.seqs’ commands in mothur. All high-quality sequences were aligned using PyNAST and dereplicated with UCLUST (Caporaso et al. 2010a) in QIIME (Quantitative Insights Into Microbial Ecology, v1.9.1)(Caporaso et al. 2010b). Finally, the Greengenes database (version 13_8)(DeSantis et al. 2006) was used as the reference database for classifying *de novo* operational taxonomic units (OTUs) that were clustered at the 97% nucleotide identity threshold. We set 0.001% as the threshold to filter the low-abundance OTUs and keep abundant OTUs, i.e., only OTUs with the read counts >0.001% of the total reads of all samples were kept for analysis(Li et al. 2015).

### Microbial diversity assessment

Microbial alpha- and beta-diversity values were determined using the QIIME(Caporaso et al. 2010b) pipeline. For alpha-diversity, rarefaction curves were drawn based on the following metrics: Observed OTUs, Chao1, Phylogenetic Diversity (PD) Whole Tree metric, and the Shannon evenness metric(Magurran 2013). For beta-diversity analysis, the final OTU table was rarefied to contain 61,088 reads per sample. Bray-Curtis, weighted and unweighted UNiFrac distance metrics (Lozupone and Knight 2005) were used to measure community similarity between samples. Microbial community clustering was arrayed by Principle Coordinate Analysis (PCoA) and visualized using Emperor(Vázquez-Baeza et al. 2013) in QIIME. The hierarchical clustering method, UPGMA (Unweighted Pair Group Method with Arithmetic Mean), was applied to cluster all water and sediment samples, and the clustering tree was visualized in FigTree (version 1.4.2)(Rambaut 2014). Permutational multivariate analysis of variance (PERMANOVA)(Anderson 2001) was performed on the Bray-Curtis distance matrix to compare differences in community structure.

### Functional profiling

PICRUSt (version 1.0.0-dev)(Langille et al. 2013) was used to make functional predictions based on the 16S rDNA dataset from each sample. For this, OTU-picking was performed on all quality-filtered sequence data using the ‘pick_closed_reference_otus.py’ command in QIIME. OTUs were clustered at the 97% nucleotide identity threshold using the Greengenes database. The OTU table was normalized using the ‘normalize_by_copy_number.py’ command. The normalized OTU table was used for functional prediction with the ‘predict_metagenomes.py’ script, and functional trait abundances were determined for each sample using the KEGG database (version 66.1, May 1, 2013)(Kang et al. 2016). Finally, the predicted functional content was collapsed to level three of the KEGG hierarchy using the ‘categorize_by_function.py’ script.

### Analysis of the relationships between physicochemical properties, antibiotics, and microbial communities

Canonical correspondence analysis (CCA) was used to identify an environmental basis for community ordination, revealing relationships between microbial communities and environmental factors(Ter Braak 1986). For this, the CCA function in R package, vegan was utilized. We utilized the ‘envfit’ function(Dawson et al. 2012, Virtanen et al. 2009) with 999 permutations to reveal significant correlations between physicochemical properties, antibiotics, and microbial communities. To further investigate correlations between environmental factors (including physicochemical properties and antibiotics) and OTUs, we applied a low-abundance filter to remove OTUs whose relative abundance did not exceed 0.01% in any sample (as previously reported by (Sunagawa et al. 2015)). Similarly, for physicochemical data and antibiotics data, the values of each variable were transformed to *z*-scores(Crocker and Algina 1986), based on which the Pearson Correlation Coeffcient between each environmental factor and each OTU was calculated. To select for significant interactions between an environmental factor and an OTU, the threshold of the r-value and the False Discovery Rate (FDR)-corrected-*p*-value of the Pearson Correlation Coeffcient was set at 0.8 and 0.05, respectively.

### Analysis of environmental drivers of microbial community composition

We noted environmental drivers of microbial community composition on the basis of (i) compositional data, which includes taxonomic composition (relative taxonomic abundances) and functional composition at KEGG module level three; (ii) physicochemical data; and (iii) antibiotics data. To pre-process compositional data, we applied a low-abundance filter to remove OTUs whose relative abundance did not exceed 0.01% in any sample and then log transformed the relative abundances. Likewise, for physicochemical and antibiotics data, the values of each variable were transformed to *z*-scores. Based on the Euclidean distances, we computed Mantel correlations between the physicochemical data and compositional data and then the antibiotics data and compositional data (9,999 permutations). We obtained the results in R (version 3.3.1) and visualized it in the Adobe Illustrator (version 16.0.0). Taxonomic composition and functional composition data were correlated to each antibiotic and physicochemical property by Mantel’s tests. The distance correlations and the statistical significance of Mantel’s r statistic corresponded to edge width and edge color, respectively(Sunagawa et al. 2015).

### Biomarker analysis

Based on their location, all water and sediment samples can be divided into two groups—impacted and less-impacted (control) groups. It is well known that the taxonomic composition of a microbial community can be impacted by local environmental variables. As a result, some bacteria might be enriched by distinctive environmental states. Linear discriminate analysis (LDA) effect size (LEfSe)(Segata et al. 2011) was used to select biomarkers in impacted and less-impacted (control) groups in water and sediment samples. Briefly, the taxa abundance table was imported into the LEfSe pipeline, and the parameters were set as follows: the alpha value for the factorial Kruskal-Wallis test(Breslow 1970) among classes and the *p*-value for the pairwise Wilcoxon test between subclasses were both chosen to be 0.05. The threshold for the logarithmic LDA score for discriminative features was set at 3.0 and 3.5 for water and sediment samples, respectively.

### Co-occurrence network analysis

To reduce sparsity, we selected water and sediment OTUs that were present in at least 50% of all water and sediment samples, respectively. We then generated separate networks for water and sediment microbial communities. The co-occurrence network was constructed using the CAVNet package(Cardona 2017) in R (as previously described by(Ma et al. 2016)). Briefly, water and sediment networks were inferred using the Spearman correlation matrix with the WGCNA package(Langfelder and Horvath 2012). In this network, co-occurring OTUs are represented by nodes and connected by edges. The network deconvolution method was utilized to distinguish direct correlation dependencies(Feizi et al. 2013). All *p*-values were corrected for multiple testing using the Benjamini and Hochberg FDR-controlling procedure(Benjamini et al. 2006). The cutoff of the FDR-corrected-*p*-value was set at 0.01. Random matrix theory-based methods were utilized to determine the cutoff of Spearman correlation coefficients for water (0.84) and sediment (0.81) networks. All network properties were calculated using the igraph package in R(Csardi and Nepusz 2006). We also utilized igraph to visualize and generate water and sediment networks. The WalkTrap community detection algorithm was used to identify modules in water and sediment networks(Pons and Latapy 2005). To study the effect of prolonged agricultural practices, we colored each node within the water and sediment network as function of its relative abundance at impacted and less-impacted (control) sites using the ‘plot_network_by_continuous_variable’ function in CAVNet.

## Declarations

### Authors’ contributions

The whole study was designed by ZW and KN. MZH, JQZ and ZW collected samples. MZH, CYC, QY and HZ performed DNA extraction and sequencing. MZH, CYZ, MD and HJL analyzed the data. MZH, CYZ, MD, HJL, JG, ZW, and KN wrote the initial draft of the manuscript. All revised the manuscript.

### Availability of data and material

All sequencing data for the 14 water samples and the 14 sediment samples were deposited into NCBI’s Sequence Read Archive (SRA) database under the Bioproject number PRJNA352457.

### Ethics approval and consent to participate

Not applicable.

### Consent for publication

Not applicable.

## Acknowledgements

Not applicable.

## References

Anderson, M.J. (2001) A new method for non-parametric multivariate analysis of variance. Austral Ecology 26(1), 32–46.

Babujia, L.C., Silva, A.P., Nakatani, A.S., Cantão, M.E., Vasconcelos, A.T.R., Visentainer, J.V and Hungria, M. (2016) Impact of long-term cropping of glyphosate-resistant transgenic soybean [Glycine max (L.) Merr.] on soil microbiome. Transgenic Research, 1–16.

Ban, X., Wu, Q., Pan, B., Du, Y. and Feng, Q. (2014) Application of Composite Water Quality Identification Index on the water quality evaluation in spatial and temporal variations: a case study in Honghu Lake, China. Environmental monitoring and assessment 186(7), 4237–4247.

Bao, S. (2005) Agricultural and chemistry analysis of soil. Agric. Press, Beijing, China.

Baquero, F., Martínez, J.-L. and Cantón, R. (2008) Antibiotics and antibiotic resistance in water environments. Current Opinion In Biotechnology 19(3), 260–265.

Beck, D.A., Kalyuzhnaya, M.G., Malfatti, S., Tringe, S.G., del Rio, T.G., Ivanova, N., Lidstrom, M.E. and Chistoserdova, L. (2013) A metagenomic insight into freshwater methane-utilizing communities and evidence for cooperation between the Methylococcaceae and the Methylophilaceae. PeerJ 1, e23.

Benjamini, Y., Krieger, A.M. and Yekutieli, D. (2006) Adaptive linear step-up procedures that control the false discovery rate. Biometrika, 491–507.

Bowles, T.M., Acosta-Martínez, V., Calderón, F. and Jackson, L.E. (2014) Soil enzyme activities, microbial communities, and carbon and nitrogen availability in organic agroecosystems across an intensively-managed agricultural landscape. Soil Biology and Biochemistry 68, 252–262.

Breslow, N. (1970) A generalized Kruskal-Wallis test for comparing K samples subject to unequal patterns of censorship. Biometrika 57(3), 579–594.

Brooks, B.W., Lazorchak, J.M., Howard, M.D., Johnson, M.V.V, Morton, S.L., Perkins, D.A., Reavie, E.D., Scott, G.I., Smith, S.A. and Steevens, J.A. (2016) Are harmful algal blooms becoming the greatest inland water quality threat to public health and aquatic ecosystems? Environmental Toxicology and Chemistry 35(1), 6–13.

Caporaso, J.G., Bittinger, K., Bushman, F.D., DeSantis, T.Z., Andersen, G.L. and Knight, R. (2010a) PyNAST: a flexible tool for aligning sequences to a template alignment. Bioinformatics 26(2), 266–267.

Caporaso, J.G., Kuczynski, J., Stombaugh, J., Bittinger, K., Bushman, F.D., Costello, E.K., Fierer, N., Pena, A.G., Goodrich, J.K. and Gordon, J.I. (2010b) QIIME allows analysis of high-throughput community sequencing data. Nature Methods 7(5), 335–336.

Cardona, C. (2017) CAVNet.

Chávez-Romero, Y., Navarro-Noya, Y.E., Reynoso-Martínez, S.C., Sarria-Guzmán, Y., Govaerts, B., Verhulst, N., Dendooven, L. and Luna-Guido, M. (2016) 16S metagenomics reveals changes in the soil bacterial community driven by soil organic C, N-fertilizer and tillage-crop residue management. Soil and Tillage Research 159, 1–8.

Chen, M., Chen, J. and Sun, F. (2008) Agricultural phosphorus flow and its environmental impacts in China. Science of The Total Environment 405(1), 140–152.

Cheng, X., Chen, X., Su, X., Zhao, H., Han, M., Bo, C., Xu, J., Bai, H. and Ning, K. (2014a) DNA extraction protocol for biological ingredient analysis of LiuWei DiHuang Wan. Genomics, Proteomics & Bioinformatics 12(3), 137–143.

Cheng, X., Su, X., Chen, X., Zhao, H., Bo, C., Xu, J., Bai, H. and Ning, K. (2014b) Biological ingredient analysis of traditional Chinese medicine preparation based on high-throughput sequencing: the story for Liuwei Dihuang Wan. Scientific Reports 4.

Cherkasov, A., Hilpert, K., Jenssen, H., Fjell, C.D., Waldbrook, M., Mullaly, S.C., Volkmer, R. and Hancock, R.E. (2008) Use of artificial intelligence in the design of small peptide antibiotics effective against a broad spectrum of highly antibiotic-resistant superbugs. ACS chemical biology 4(1), 65–74.

Crocker, L. and Algina, J. (1986) Introduction to classical and modern test theory, ERIC.

Csardi, G. and Nepusz, T. (2006) The igraph software package for complex network research. InterJournal, Complex Systems 1695(5), 1–9.

Dawson, K.S., Strąpoć, D., Huizinga, B., Lidstrom, U., Ashby, M. and Macalady, J.L. (2012) Quantitative fluorescence in situ hybridization analysis of microbial consortia from a biogenic gas field in Alaska’s Cook Inlet Basin. Applied and Environmental Microbiology 78(10), 3599–3605.

Dennis, P.G., Seymour, J., Kumbun, K. and Tyson, G.W. (2013) Diverse populations of lake water bacteria exhibit chemotaxis towards inorganic nutrients. The ISME Journal 7(8), 1661–1664.

DeSantis, T.Z., Hugenholtz, P., Larsen, N., Rojas, M., Brodie, E.L., Keller, K., Huber, T., Dalevi, D., Hu, P. and Andersen, G.L. (2006) Greengenes, a chimera-checked 16S rRNA gene database and workbench compatible with ARB. Applied and Environmental Microbiology. 72(7), 5069–5072.

Divyashree, M.S., Rastogi, N.K. and Shamala, T.R. (2009) A simple kinetic model for growth and biosynthesis of polyhydroxyalkanoate in Bacillus flexus. New Biotechnology 26(1), 92–98.

Federation, W.E. and Association, A.P.H. (2005) Standard methods for the examination of water and wastewater. American Public Health Association (APHA): Washington, DC, USA.

Feizi, S., Marbach, D., Médard, M. and Kellis, M. (2013) Network deconvolution as a general method to distinguish direct dependencies in networks. Nature biotechnology 31(8), 726–733.

Gechemba, O.R., Budambula, N., Makonde, H.M., Julius, M. and Matiru, V.N. (2015) Potentially beneficial rhizobacteria associated with banana plants in Juja, Kenya. Journal of Biodiversity and Environmental Sciences 7(2), 181–188.

Geist, J. and Hawkins, S.J. (2016) Habitat recovery and restoration in aquatic ecosystems: current progress and future challenges. Aquatic Conservation: Marine and Freshwater Ecosystems.

Guo, D., Guan, L., Zhang, C., Wang, X. and Shan, L. (2013) UV Mutagenesis Breeding of Bacillus flexus Highly Degrading Organic Nitrogen. Guizhou Agricultural Sciences 11, 028.

Han, M., Gong, Y., Zhou, C., Zhang, J., Wang, Z. and Ning, K. (2016) Comparison and Interpretation of Taxonomical Structure of Bacterial Communities in Two Types of Lakes on Yun-Gui plateau of China. Sci Rep 6.

Horppila, J. and Nurminen, L. (2003) Effects of submerged macrophytes on sediment resuspension and internal phosphorus loading in Lake Hiidenvesi (southern Finland). Water research 37(18), 4468–4474.

Kang, C., Zhang, Y., Zhu, X., Liu, K., Wang, X., Chen, M., Wang, J., Chen, H., Hui, S. and Huang, L. (2016) Healthy subjects differentially respond to dietary capsaicin correlating with specific gut enterotypes. The Journal of Clinical Endocrinology and Metabolism. 101(12), 4681–4689.

Langfelder, P. and Horvath, S. (2012) Fast R functions for robust correlations and hierarchical clustering. Journal of statistical software 46(11).

Langille, M.G., Zaneveld, J., Caporaso, J.G., McDonald, D., Knights, D., Reyes, J.A., Clemente, J.C., Burkepile, D.E., Thurber, R.L.V. and Knight, R. (2013) Predictive functional profiling of microbial communities using 16S rRNA marker gene sequences. Nature Biotechnology. 31(9), 814–821.

Lee, L.S., Carmosini, N., Sassman, S.A., Dion, H.M. and Sepulveda, M.S. (2007) Agricultural contributions of antimicrobials and hormones on soil and water quality. Advances In Agronomy 93, 1–68.

Li, J., Zhang, J., Liu, L., Fan, Y., Li, L., Yang, Y., Lu, Z. and Zhang, X. (2015) Annual periodicity in planktonic bacterial and archaeal community composition of eutrophic Lake Taihu. Scientific Reports 5.

Lindström, E.S., Kamst-Van Agterveld, M.P. and Zwart, G. (2005) Distribution of typical freshwater bacterial groups is associated with pH, temperature, and lake water retention time. Applied and Environmental Microbiology 71(12), 8201–8206.

Lozupone, C. and Knight, R. (2005) UniFrac: a new phylogenetic method for comparing microbial communities. Applied and Environmental Microbiology. 71(12), 8228–8235.

Ma, B., Wang, H., Dsouza, M., Lou, J., He, Y., Dai, Z., Brookes, P.C., Xu, J. and Gilbert, J.A. (2016) Geographic patterns of co-occurrence network topological features for soil microbiota at continental scale in eastern China. The ISME Journal.

Magurran, A.E. (2013) Measuring biological diversity, John Wiley & Sons.

Nielsen, P.H., Kragelund, C., Seviour, R.J. and Nielsen, J.L. (2009) Identity and ecophysiology of filamentous bacteria in activated sludge. FEMS microbiology reviews 33(6), 969–998.

O’neil, J., Davis, T.W., Burford, M.A. and Gobler, C. (2012) The rise of harmful cyanobacteria blooms: the potential roles of eutrophication and climate change. Harmful Algae 14, 313–334.

Pei, R., Kim, S.-C., Carlson, K.H. and Pruden, A. (2006) Effect of river landscape on the sediment concentrations of antibiotics and corresponding antibiotic resistance genes (ARG). Water research 40(12), 2427–2435.

Pesaro, M. and Widmer, F. (2006) Identification and specific detection of a novel Pseudomonadaceae cluster associated with soils from winter wheat plots of a long-term agricultural field experiment. Applied And Environmental Microbiology 72(1), 37–43.

Pons, P. and Latapy, M. (2005) Computing communities in large networks using random walks. International Symposium on Computer and Information Sciences, 284–293.

Porebski, S., Bailey, L.G. and Baum, B.R. (1997) Modification of a CTAB DNA extraction protocol for plants containing high polysaccharide and polyphenol components. Plant molecular biology reporter 15(1), 8–15.

Quast, C., Pruesse, E., Yilmaz, P., Gerken, J., Schweer, T., Yarza, P., Peplies, J. and Glöckner, F.O. (2013) The SILVA ribosomal RNA gene database project: improved data processing and web-based tools. Nucleic Acids Research. 41(D1), D590–D596.

Rambaut, A. (2014) FigTree v1. 4.2. Edinburgh: University of Edinburgh.

Sand-jensen, K. (1998) Influence of submerged macrophytes on sediment composition and near-bed flow in lowland streams. Freshwater biology 39(4), 663–679.

Schloss, P.D., Westcott, S.L., Ryabin, T., Hall, J.R., Hartmann, M., Hollister, E.B., Lesniewski, R.A., Oakley, B.B., Parks, D.H. and Robinson, C.J. (2009) Introducing mothur: open-source, platform-independent, community-supported software for describing and comparing microbial communities. Applied and Environmental Microbiology. 75(23), 7537–7541.

Segata, N., Izard, J., Waldron, L., Gevers, D., Miropolsky, L., Garrett, W.S. and Huttenhower, C. (2011) Metagenomic biomarker discovery and explanation. Genome Biology 12(6), R60.

Sengupta, S., Chattopadhyay, M.K. and Grossart, H.-P. (2013) The multifaceted roles of antibiotics and antibiotic resistance in nature. Frontiers in Microbiology 4.

Shao, K., Gao, G., Wang, Y., Tang, X. and Qin, B. (2013) Vertical diversity of sediment bacterial communities in two different trophic states of the eutrophic Lake Taihu, China. Journal of Environmental Sciences. 25(6), 1186–1194.

Sunagawa, S., Coelho, L.P., Chaffron, S., Kultima, J.R., Labadie, K., Salazar, G., Djahanschiri, B., Zeller, G., Mende, D.R. and Alberti, A. (2015) Structure and function of the global ocean microbiome. Science 348(6237), 1261359.

Sundaramanickam, A., Kumar, P.S., Kumaresan, S. and Balasubramanian, T. (2015) Isolation and molecular characterization of multidrug-resistant halophilic bacteria from shrimp farm effluents of Parangipettai coastal waters. Environmental Science and Pollution Research 22(15), 11700–11707.

Takamura N, K.Y., Fukushima M, Nakagawa M, Kim, B. H. (2003) Effects of aquatic macrophytes on water quality and phytoplankton communities in shallow lakes. Ecological Research 18(4), 381–395.

Ter Braak, C.J. (1986) Canonical correspondence analysis: a new eigenvector technique for multivariate direct gradient analysis. Ecology 67(5), 1167–1179.

Vázquez-Baeza, Y., Pirrung, M., Gonzalez, A. and Knight, R. (2013) EMPeror: a tool for visualizing high-throughput microbial community data. Gigascience 2(1), 1.

Verdonschot, R., Keizer-vlek, H.E. and Verdonschot, P.F. (2011) Biodiversity value of agricultural drainage ditches: a comparative analysis of the aquatic invertebrate fauna of ditches and small lakes. Aquatic Conservation: Marine and Freshwater Ecosystems 21(7), 715–727.

Virtanen, R., Ilmonen, J., Paasivirta, L. and Muotka, T. (2009) Community concordance between bryophyte and insect assemblages in boreal springs: a broad-scale study in isolated habitats. Freshwater Biology 54(8), 1651–1662.

Wang, H., Sangwan, N., Li, H.-Y., Su, J.-Q., Oyang, W.-Y., Zhang, Z.-J., Gilbert, J.A., Zhu, Y.-G., Ping, F. and Zhang, H.-L. (2017a) The antibiotic resistome of swine manure is significantly altered by association with the Musca domestica larvae gut microbiome. The ISME Journal 11(1), 100–111.

Wang, X. and Zhao, H. (2013) Isolation and Characterization of a Bacillus flexus Strain Used in Alkaline Wastewater Treatment. Advanced Materials Research 750, 1381–1384.

Wang, Z., Du, Y., Yang, C., Liu, X., Zhang, J., Li, E., Zhang, Q. and Wang, X. (2017b) Occurrence and ecological hazard assessment of selected antibiotics in the surface waters in and around Lake Honghu, China. Science of The Total Environment 609(Supplement C), 1423–1432.

Williams, C.J., Frost, P.C., Morales-Williams, A.M., Larson, J.H., Richardson, W.B., Chiandet, A.S. and Xenopoulos, M.A. (2016) Human activities cause distinct dissolved organic matter composition across freshwater ecosystems. Global change biology 22(2), 613–626.

Williams, P., Whitfield, M., Biggs, J., Bray, S., Fox, G., Nicolet, P. and Sear, D. (2004) Comparative biodiversity of rivers, streams, ditches and ponds in an agricultural landscape in Southern England. Biological conservation 115(2), 329–341.

Wood, S.A., Maier, M.Y., Puddick, J., Pochon, X., Zaiko, A., Dietrich, D.R. and Hamilton, D.P. (2017) Trophic state and geographic gradients influence planktonic cyanobacterial diversity and distribution in New Zealand lakes. FEMS Microbiology Ecology 93(2), fiw234.

Xu, H., Paerl, H.W., Qin, B., Zhu, G. and Gao, G. (2010) Nitrogen and phosphorus inputs control phytoplankton growth in eutrophic Lake Taihu, China. Limnology and Oceanography 55(1), 420.

Yang, Y., Cao, X., Lin, H. and Wang, J. (2016) Antibiotics and Antibiotic Resistance Genes in Sediment of Honghu Lake and East Dongting Lake, China. Microbial Ecology 72(4), 791–801.

Ye, W., Liu, X., Lin, S., Tan, J., Pan, J., Li, D. and Yang, H. (2009) The vertical distribution of bacterial and archaeal communities in the water and sediment of Lake Taihu. FEMS Microbiology Ecology 70(2), 263–276.

Zhang, T., Ban, X., Wang, X., Cai, X., Li, E., Wang, Z., Yang, C., Zhang, Q. and Lu, X. (2017) Analysis of nutrient transport and ecological response in Honghu Lake, China by using a mathematical model. Science of The Total Environment 575, 418–428.

Zhang, X. (1998) On the estimation of biomass of submerged vegetation using Landsat thematic mapper (TM) imagery: a case study of the Honghu Lake, PR China. International Journal Of Remote Sensing 19(1), 11–20.

